# Remote-controlled insect navigation using plasmonic nanotattoos

**DOI:** 10.1101/2020.02.10.942540

**Authors:** Sirimuvva Tadepalli, Sisi Cao, Debajit Saha, Keng-Ku Liu, Alex Chen, Sang hyun Bae, Baranidharan Raman, Srikanth Singamaneni

## Abstract

Developing insect cyborgs by integrating external components (optical, electrical or mechanical) with biological counterparts has a potential to offer elegant solutions for complex engineering problems.^1^ A key limiting step in the development of such biorobots arises at the nano-bio interface, *i.e.* between the organism and the nano implant that offers remote controllability.^1,2^ Often, invasive procedures are necessary that tend to severely compromise the navigation capabilities as well as the longevity of such biorobots. Therefore, we sought to develop a non-invasive solution using plasmonic nanostructures that can be photoexcited to generate heat with spatial and temporal control. We designed a ‘nanotattoo’ using silk that can interface the plasmonic nanostructures with a biological tissue. Our results reveal that both structural and functional integrity of the biological tissues such as insect antenna, compound eyes and wings were preserved after the attachment of the nanotattoo. Finally, we demonstrate that insects with the plasmonic nanotattoos can be remote controlled using light and integrated with functional recognition elements to detect the chemical environment in the region of interest. In sum, we believe that the proposed technology will play a crucial role in the emerging fields of biorobotics and other nano-bio applications.

---

Insects often provide compact solutions to several complex engineering problems and serve as an inspiration for developing insect-like robots that attempt to mimic those solutions.^3^ However, the ability of such robots’ pale when compared to the versatility of the biological organism on which it is modeled upon. Could ‘bio-robots’ that directly take advantage of the biological capabilities be developed by directly controlling the insects? We sought to examine these issues in this study using locusts (*Schistocerca americana*) as a model organism.

We began by designing a communication system that would allow us to directly send a control signal and guide locust’s behavioral responses. Prior work in this regard have investigated the use of electrical stimulation, whereby electrodes are surgically implanted into different parts of the insect body. ^4^ While this approach is feasible, it requires invasive surgical manipulations that could reduce the locomotor capabilities of the insects. Further, they would require the insects to carry a payload comprising at a bare minimum a receiver powered by a battery to deliver the electrical stimulation. The additional weight can once again reduce the ability of the insects to move freely.

Therefore, to develop a non-invasive, powerless approach to control locust navigation, we designed a ‘plasmonic nanotattoo.’ The nanotattoo is comprised of two components: gold nanorods (AuNRs) that act as a photothermal transducers and an ultrathin silk fibroin interfacial film. We sought out to take advantage of gold nanorods as nanotransducers that convert light at specific wavelength into localized heat (Fig. 1A-C).^5^ However, we found that these AuNR by themselves do not readily adsorb onto biological tissue surfaces. Treatment of the biological surfaces to make them conducive for AuNR adsorption resulted in compromising the structure or functionality or both of the target tissue. Therefore, to integrate these two non-compatible components, we examined the use of a silk film as an interfacial material. In addition to its excellent mechanical properties, we found that the silk film conforms to arbitrary spatial patterns owing to its nanoscale thickness (Fig. 1D-F). Importantly, we found gold nanorods readily adsorbed onto the silk film and the optical properties were preserved upon adsorption onto the silk film (Fig. 1C).

**Figure 1.**
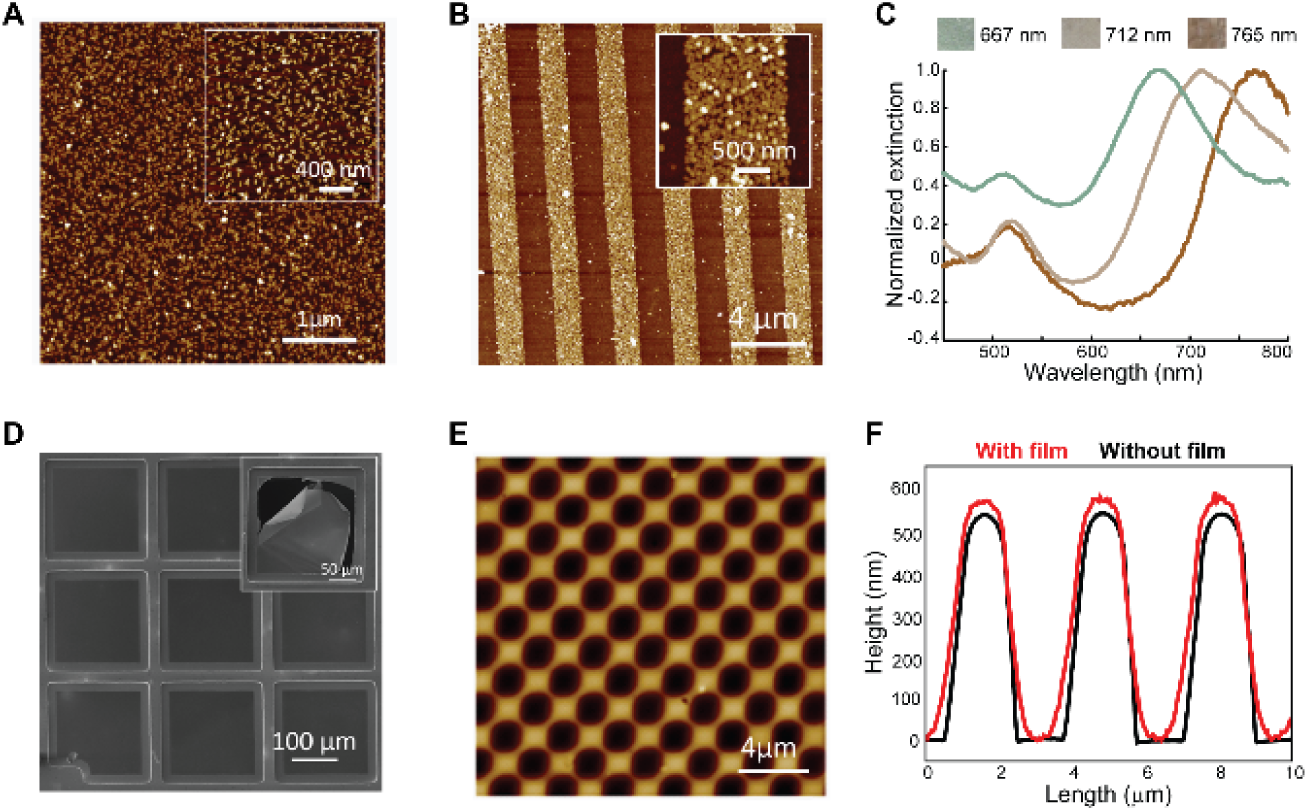
(A) AFM image of the silk film after AuNR adsorption showing the uniformly adsorbed AuNR. Z-scale: 35 nm. (B) Silk film with patterned AuNRs on the surface showing the ability control the spatial distribution of the AuNRs. Inset showing the magnified image of the AuNR stripe. Z-scale: 50 nm. (C) Plasmonic silk films with AuNRs of different LSPR wavelengths. (D) SEM image of the silk film transferred on to a TEM grid. Inset shows the image of an intentionally broken region. (E) Conformality of the silk film on a patterned substrate. Z-scale: 900 nm. (F) AFM height profile of the pattern with film showing that the film is conformal on the pattern.

Next, we examined if the gold-studded silk film (nanotattoo) can adsorb onto biological surfaces without significantly impacting the structure and function of the target tissue. The ability of these ultrathin films to cover and conform to a surface with complex topography is critical to exploit a number of unique optical properties of plasmonic nanostructures embedded on them. Hence, we examined whether the nanotatto can be transferred on biological surfaces with varied physiochemical properties such as insect antenna and compound eyes (Fig. 2A-C). We found that indeed the tattoos were able to faithfully conform to the structure and morphology of these varied biological tissues. These results therefore confirm that silk acts as an excellent interfacial material at nano-bio interfaces.

**Figure 2.**
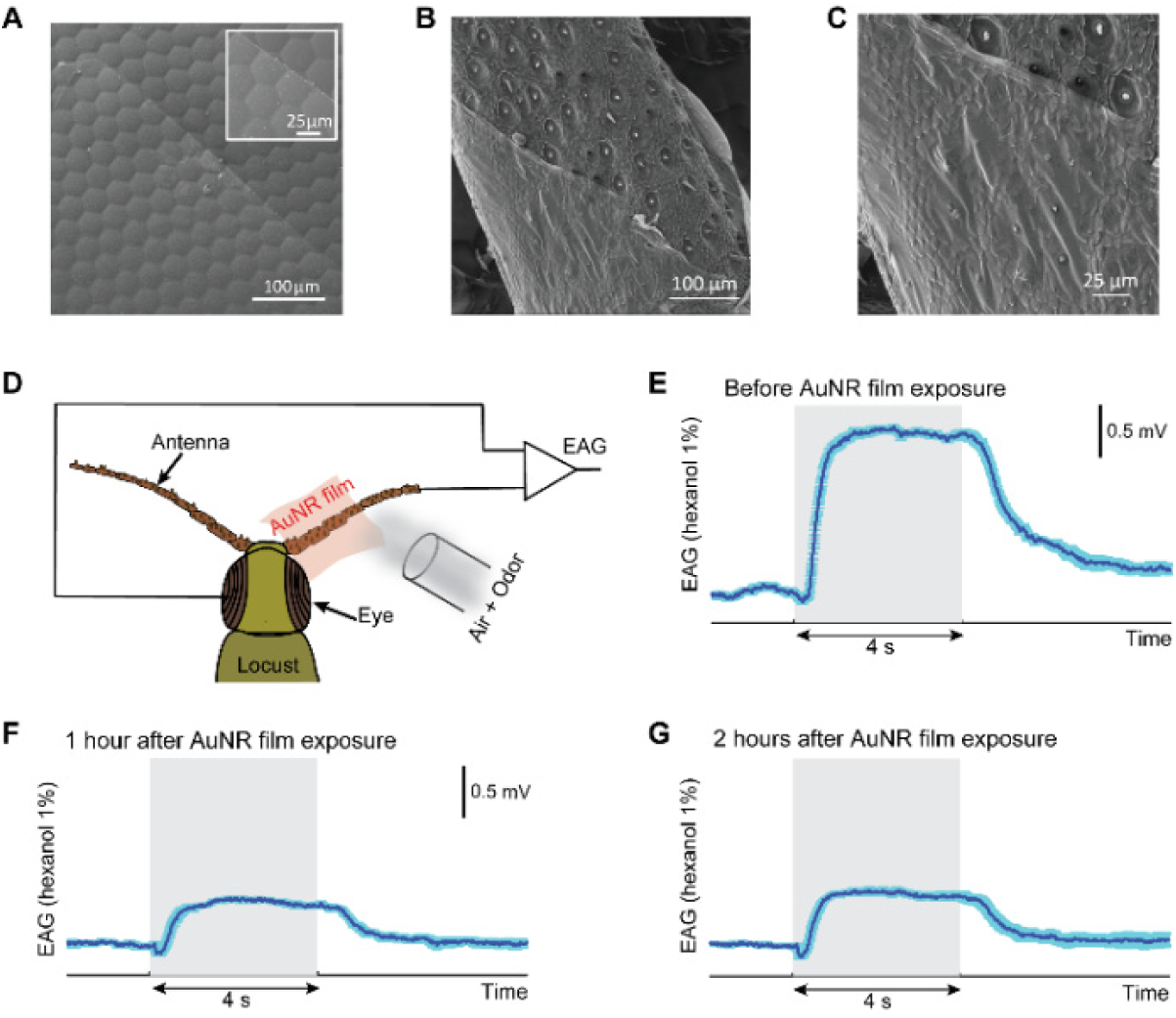
SEM images of the film transferred on to a complex biological surface (A) Compound Eye (B, C) antenna. (D) Schematic depicting the EAG measurement setup. EAG measurement on locust antenna with 1% hexanol odor puff (E) before AuNR film (F) 1 hour after AuNR film and (G) 2 hours after AuNR film.

Are the functionalities of these biological surfaces retained after the application of the silk film? We examined the excitability of the insect antenna before and after the application of a silk film (Fig. 2D-G). We found that the electroantennogram that represent the total activity of olfactory and non-olfactory neurons housed in insect antenna after application of the silk film reduced in response amplitude during odor exposure. This probably was due to the silk film covering a substantial portion of the antennal surface area and thereby preventing odor molecules from reaching the sensory hairs covered by them. However, the presence of the silk film did not negatively impact the function of the antennal tissue surrounding it even several hours after its application (Fig. 2G). These results confirm that the silk film provides an excellent interface between the synthesized nanomaterials with special transduction properties and biological surfaces with special structure and functional capabilities.

Given that the gold nanorods transduce light to heat that could potentially repel insects, we wondered if a patch of silk film tattooed onto the locust could help control its locomotion. To test this, we transferred our ‘nanotattoo’ to locust wings (Fig. 3A). When excited with light (laser with a wavelength of 808 nm), we found that the temperature on the tattooed wing increased by 20°C within 20 seconds (Fig. 3B, C). Note that the heating was local to the region bearing the nanotattoo. As a control, direct stimulation of a non-tattooed wing resulted only in modest increase in the temperature (Fig. 3B, C). We found that tattooed locusts restrained in a holding station rapidly moved out when irradiated with laser compared to the control case (Fig. 3D-F). These results taken together provide a proof-of-concept demonstration that the tattooed locusts can be optically controlled.

**Figure 3:**
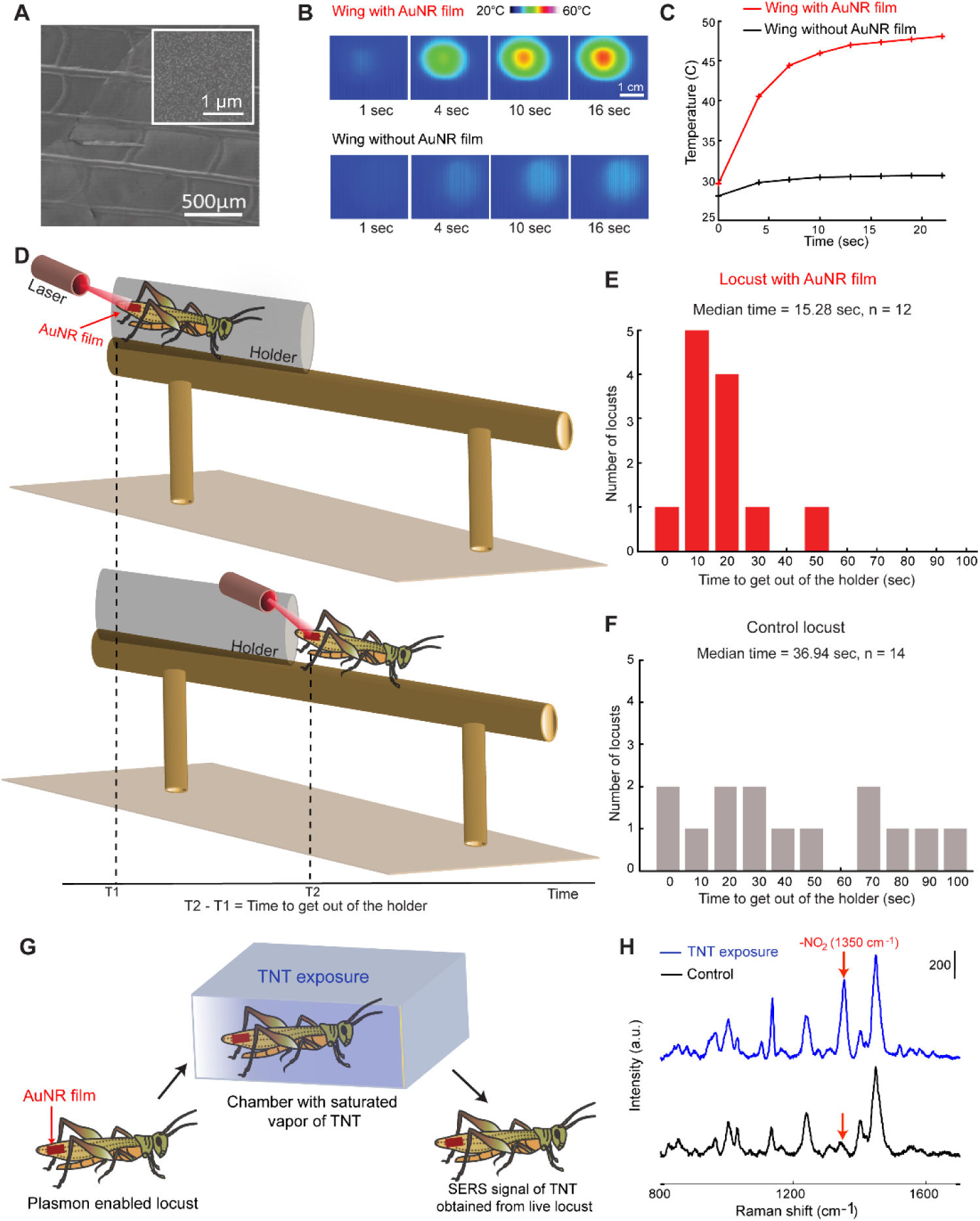
SEM image of the silk film on the wing on the locust wing (Inset: Image at high magnification) (B) Temperature profile showing the raise in temperature over 20 seconds of exposure to 808nm laser at a power of 400mW/cm^2^.(C) IR camera images collected on a live locust wing with and without plasmonic patch after exposure to 808nm laser at a power of 400mW/cm^2^. (D) Speed assay setup with locust movement. Blind test of speed assay for locust behavioral control on (E) locust with AuNR film (F) control locust. (G) Schematic showing the experimental setup to measure TNT vapors on a live locust (H) Raman spectra collected on a live locust after exposure to TNT vapors (Units: counts per second).

While there are several potential applications for an insect-based biorobot, we examined if the tattooed locusts can be used to detect the presence or absence of chemical targets in regions of interest. We explored whether the nanotatttoos can be tuned to particularly look for certain target molecules, such as trinitro toluene (TNT), a commonly used explosive. To achieve this goal, we customized the nanorods for this problem by functionalizing AuNR with peptides that selectively bind to TNT molecules. To illustrate this, we steered a locust with nanotattoo through a chamber containing TNT vapors (Fig. 3G, H). Subsequent surface enhanced Raman scattering (SERS) spectra from the locust tattoo exposed to TNT vapors indicated peaks at 1349 cm^-1^ corresponding -NO_2_ stretching of TNT. These results taken together demonstrate a simple approach to realize insect-based robot for explosive sensing.

## Discussion

There has been a phenomenal progress in the design and synthesis of organic and inorganic nanostructures with great control over the size, shape, composition, surface functionality and properties that are unique at the nanoscale.^6-8^ However, interfacing these functional nanostructures with micro- and macroscopic physical and biological systems remains a fundamental challenge. More often than not, these biological systems require a tight control over environmental factors such as temperature, pH, and solvent environment, which limits the available processing window. Furthermore, any processing of the biological tissue to make them amenable to conjugation with the nanomaterials may not preserve their structural and/or functional integrity ^9,10^. Therefore, there remains a fundamental need to develop methods that allow facile interfacing of functional nanostructures with biological systems.

One possible solution to this problem is through the use of interfacial biopolymers such as silk that has excellent mechanical properties (e.g., strength, toughness), biocompatibility and biodegradability.^11,12^ Due to its hierarchical multi-domain morphology, silk fibers are among the strongest biopolymeric materials with a unique combination of high modulus and high elongation at failure.^13^ The rich chemical functionality and the mechanical robustness of nanoscale silk fibroin films makes them ideal for bio-nano interfacial applications.^14^ Our study is the first to demonstrate that silk films can be used as an interfacial material that can seamlessly integrate tailor-made nano-transducers with biological tissue targets, and whether this could be achieved *in vivo*.

We designed a bioplasmonic silk tattoo patterned with nanoscale heating elements that readily adsorb to complex biological surfaces such as an insect antenna, compound eyes and wings. We found that these silk tattoo’s preserves the functionality of both the nanoheaters and the biological tissue. Finally, our results demonstrate that tattooed locust can be manipulated to locomote using an external trigger. In sum, our results provide a proof-of-concept demonstration of a biorobot capable of sensing explosive targets in regions of interest.

## Acknowledgement

This research was supported by McDonnell Center for Systems Neuroscience grant and Office of Naval Research grant (N000141612426 and N000141210089).

## Author contributions

B.R. and S. S. conceived the study and designed the experiments. S.T. and S.C. synthesized, systematically characterized and tested the nanorods and the silk film. K-K.L. characterized the silk film on various biological surfaces using SEM. D.S., performed the electrophysiological recordings. S.T. D.S. and A.C. conducted the behavioral experiments and analyzed the data. S.T. and S-H.B. performed the TNT experiments and took the subsequent Raman spectra measurements. S.T. and D.S. wrote the first draft of the paper, B.R. and S.S. revised the manuscript incorporating feedback from all authors. B.R. and S.S. provided overall supervision. S.C. and D.S. contributed equally to this work.

## METHODS

### Materials

Sodium carbonate, lithium bromide, polystyrene (M_w_ = 192,000), toluene, acetone, methanol, sodium borohydride, cetyltrimethylammonium bromide, gold chloride trihydrate, silver nitrate, L-ascorbic acid and hydrochloric acid were obtained from Sigma-Aldrich. Silk cocoons were purchased from Mulberry farms.

### Processing of silk

Silk fibroin was extracted from *Bombyx mori* silkworm cocoons as previously reported. ^15^ Briefly, silk cocoons were boiled for 30 minutes in an aqueous solution of sodium carbonate (Na_2_CO_3_, 0.02M) and then rinsed thoroughly with nanopure water (18.2 MΩ-cm) to remove sericin proteins. The extracted silk fibroin was dissolved in lithium bromide (LiBr, 9.3 M) solution at 60°C for 4 h. The solution was dialyzed against nanopure water at room temperature for 2 days to remove the LiBr salts. The dialysate was centrifuged twice at 4°C at 9000 rpm for 10 min, to remove impurities and the aggregates that occur during dialysis. The dialysate solution was stored at 4°C until prior to use.

### Fabrication of a free-standing silk film

For the fabrication of a free-standing silk film, a sacrificial polystyrene (PS) layer was deposited on the piranha-cleaned silicon substrate by spin coating from 2% (w/v) PS solution in toluene at 3000 rpm for 30 seconds, followed by the deposition of 2% (w/v) aqueous silk solution on top of the PS layer with the speed of 3000 rpm for 30 seconds. The substrate was then immersed in methanol solution for 3 minutes to induce β-sheet formation followed by natural drying. In order to release the silk film, the substrate was immersed in toluene solution to remove the PS sacrificial layer. The film is then transferred to water to form a free-standing film at the interface of water and air. The procedure for the film fabrication and transfer is shown in the Extended Figure 1.

### Synthesis of gold nanorods (AuNR)

AuNRs were synthesized using a previously reported seed-mediated approach.^16^ Seed solution was prepared by adding 0.6 ml of freshly prepared ice-cold sodium borohydride (NaBH_4_, 10 mM) aqueous solution into a mixture of 9.75 ml of cetyltrimethylammonium bromide (CTAB, 0.1 M) and 0.25 ml of gold chloride trihydrate (HAuCl_4_, 10 mM) solution under vigorous stirring at room temperature (10 min, 8000 rpm). The color of the seed solution changed immediately from yellow to brown after NaBH_4_ addition. Growth solution was prepared by mixing 95 ml of CTAB (0.1 M), 5.0 ml of HAuCl_4_ (10 mM), 1.0 ml of silver nitrate (10 mM) and 1.1 ml of ascorbic acid (0.1 M) in order, followed by gentle shaking. To the resulting colorless growth solution, 0.24 ml of freshly prepared seed solution was added. After overnight incubation, AuNRs solution was centrifuged twice at 10000 rpm for 10 min to remove excess reactants and dispersed in nanopure water (18.2 MΩ-cm). For the immobilization of AuNRs on the silk film, the substrate with silk film was incubated in a 3 ml solution containing twice centrifuged AuNRs (extinction 2.0) and 5 µl of 1 M HCl for 48 hours.

### Peptide Conjugation

TNT binding peptide has been identified by phage display in a recent report.^17^ We attached three glycines and one cysteine at the N-terminus of the peptide sequence (WHWQRPLMPVSI) to impart conformational flexibility to the peptide and facilitate the conjugation through gold-thiol interaction of the cysteine at the N-terminus. The silk films with AuNRs were incubated with an aqueous peptide solution (1mM) for 12 hours to conjugate the peptides with the AuNRs on the silk film.

## Supporting Information

**Extended Figure 1:**
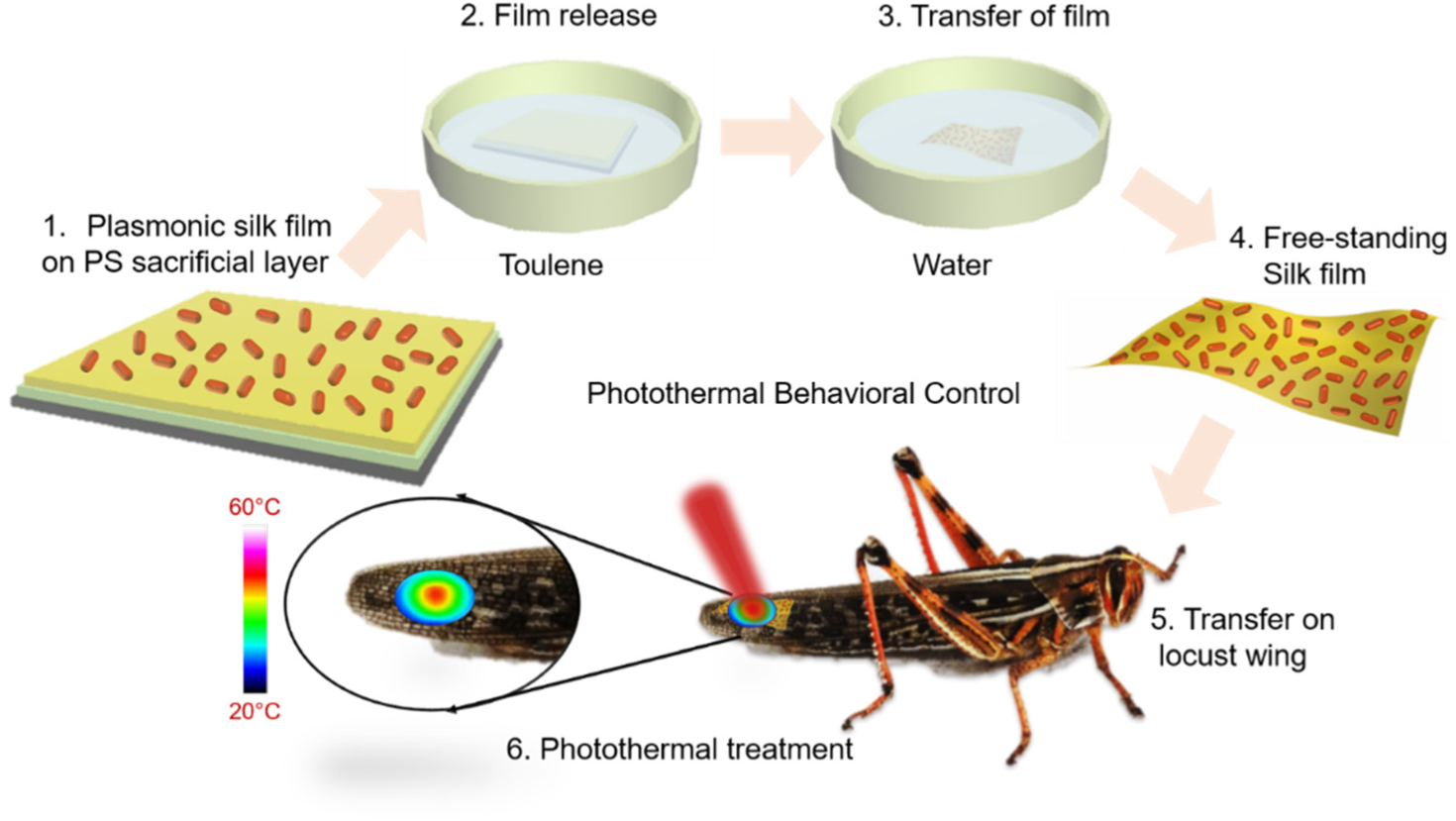
Schematic showing the concept of photothermal transduction of plasmonic silk film on a live locust wing.

**Extended Figure 2:**
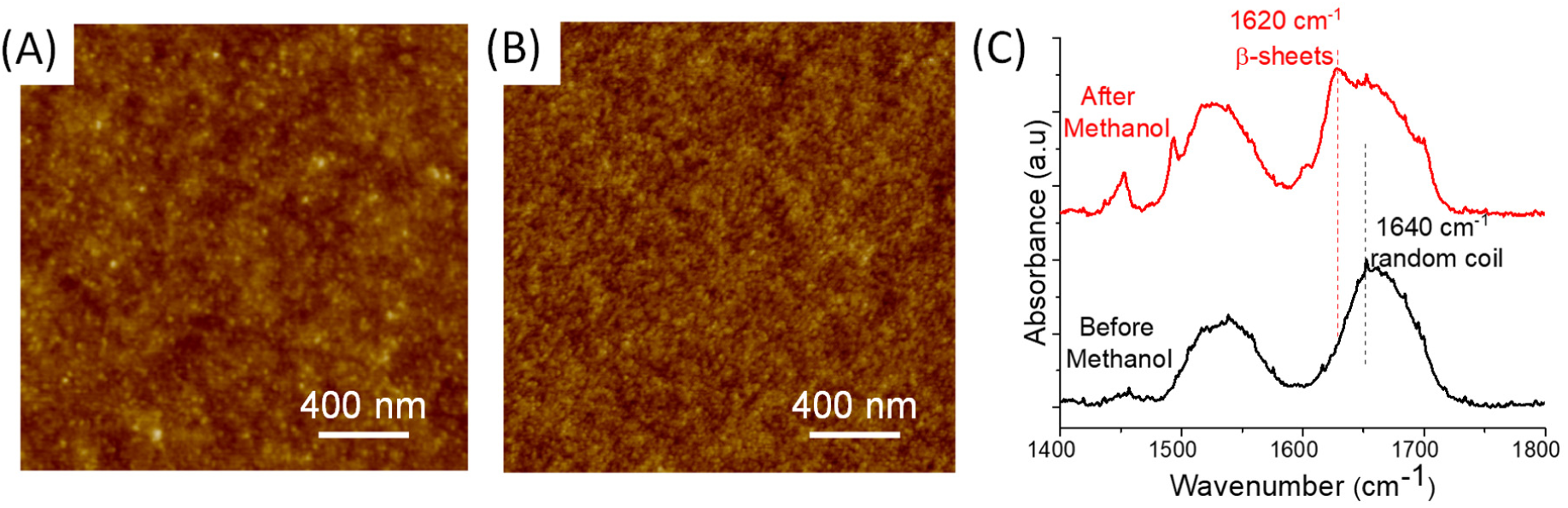
AFM image of the morphology of the silk film (A) before methanol treatment and (B) after methanol treatment. Z-scale: 10nm. (C) IR Spectra of the silk film before and after methanol treatment showing the change in the conformation.

### Characterization of a water-insoluble silk film

The silk film exposed to methanol results in a conformational change of regenerated silk fibroin with a significant increase in β-sheet making it a water insoluble film. ^1^ The *as deposited* silk film exhibited a smooth featureless morphology with a RMS roughness of 3.5 nm over 1×1 μm^2^ area. After methanol treatment, the film exhibited a grainy morphology with a RMS roughness of 5.6 nm over a 1×1 μm^2^ area (Figure 2A). The conformational change in the silk film with methanol treatment was investigated using Fourier transform infrared (FTIR) spectroscopy. The FTIR spectra of the silk films comprised of amide I (1700-1600 cm^-1^) and amide II (1600-1500 cm^-1^) corresponding to the protein backbone.^2^ After methanol treatment, the peaks in the amide II region (1648-1654 cm^-1^), corresponding to silk I conformation, shifted to (1610-1630 cm^-1^) indicating the transformation to silk II conformation (Figure 2B).^2,3^ This conformational change associated with methanol treatment is critical in the design of a water insoluble free-standing plasmonic film.

**Extended Figure 3:**
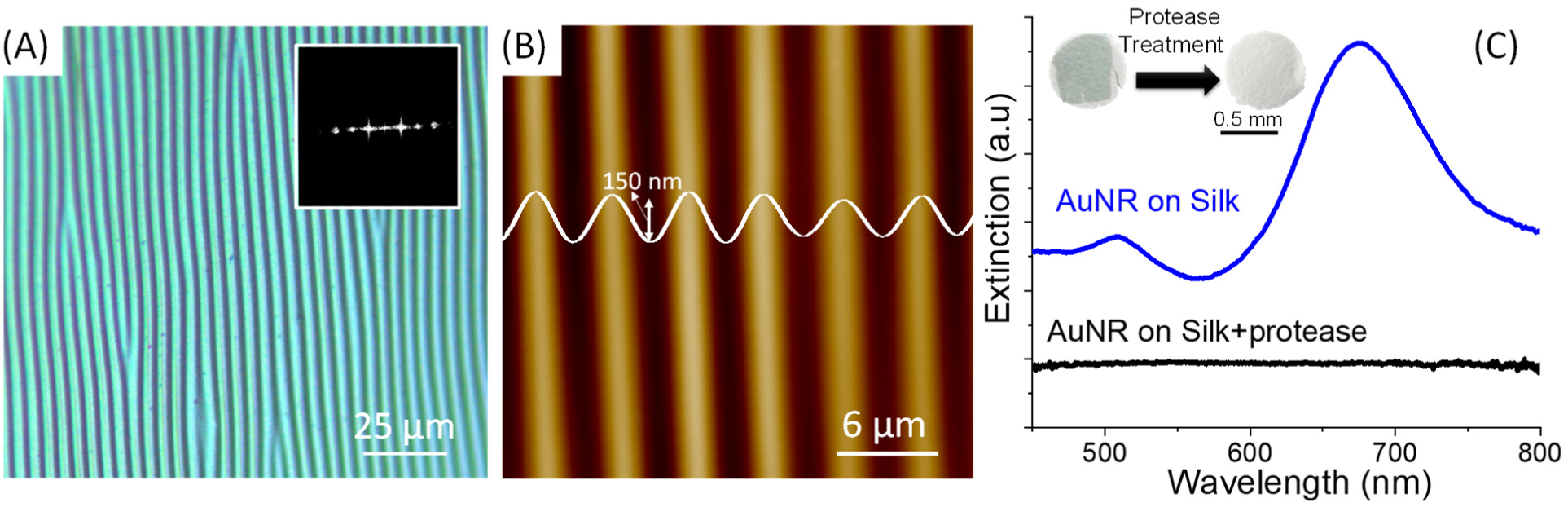
(A) Optical micrograph of buckling of the silk film. Inset shows the FFT of the buckling pattern (C) AFM topography of the buckling pattern of the free-standing film. (D) Protease treatment of silk film on paper showing the degradation of the silk film and the loss of the optical properties.

### Probing the elastic modulus of the silk film

In order to probe the elastic modulus, the free-standing film was transferred onto a compliant poly(dimethylsiloxane) (PDMS) substrate. The elastic modulus of plasmonically-active silk film was determined using strain-induced elastic buckling instability for mechanical measurements. ^4^ Briefly, due to the compressive stresses, periodic buckling patterns are formed spontaneously to minimize the strain energy at compressive stress above a certain threshold. ^5^ The buckling wavelength is given by: ^6^

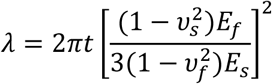

Where *λ* is the wavelength of the periodic buckling pattern, *E*_*f*_ and *ν*_*f*_ are the elastic modulus and the Poisson’s ratio of the film, and *E*_*s*_ and *ν*_*s*_ are the elastic modulus and the Poisson’s ratio of the compliant substrate and *t* is the thickness of the film.

Compression of the compliant PDMS substrate with free-standing silk film resulted in a uniform buckling of the plasmonically-active silk film. Optical images revealed large regions of periodic wrinkles with a uniform spacing (Extended Figure 3A). Fast Fourier transform (FFT) of the optical images was employed to determine the wavelength of the buckles to be 4 ± 0.2 μm (inset of Extended Figure 3A). AFM imaging of the buckled surface further confirmed the periodicity of the buckling patterns and revealed amplitude of the buckles to be ∼150 nm (Extended Figure 3B). The corresponding Young’s modulus of the silk film with GNRs was calculated to be 5.2 ± 0.3 GPa. The elastic modulus agrees with the previously reported ultrathin free-standing silk films obatined using layer-by-layer silk-on-silk assembly.^7^ Presence of GNRs results in only a modest increase in the elastic modulus compared to the pristine film (4.5 ± 0.3 GPa) due to their adsorption on the surface as opposed to being dispersed within the silk matrix. The excellent mechanical stability combined with low flexural rigidity of the nanoscale silk film is critical to transfer to complex surfaces without fracture.

### Probing the biodegradability of the silk film

In order to test the biodegradability to the film, we subjected the film transferred onto a cellulose paper surface to proteolytic digestion. Owing to the white background of the cellulose paper, the transferred film with GNR could be easily visualized under ambient light. Extinction spectra from the paper surface also revealed the LSPR bands corresponding to the GNR. Upon exposure to 1 mg/ml of protease overnight, the silk film completely disappeared (Extended Figure 3C). Further, the extinction spectrum obtained from paper substrate after subjecting the silk film to proteolytic digestion did not exhibit discernable LSPR bands suggesting that the GNRs dispersed into the solution during proteolysis. Incubation of paper substrates with plasmonically-active silk film to aqueous medium without protease did not affect the silk film, confirming that the biodegradation of the film is due to the presence of protease.

### Electroantennogram (EAG) recordings

EAG recordings were made from intact locust antenna before and after silk film exposure. A young-adult locust (Schistocerca americana) raised in a crowded colony was used for these experiments. For these recordings, 3 to 4 distal segments of the locust antenna were cut and an Ag/AgCl wire was inserted into the antenna. The counter electrode, another Ag/AgCl wire, was inserted into the contralateral eye. EAG signals were amplified using a DC amplifier (Brownlee Precision) and acquired at 15 kHz sampling rate (PCI-MIO-16E-4 DAQ cards; National Instruments).

### Behavioral experiments

Young-adult locusts (post-fifth instar) of either sex were used in these experiments. A silk film studded with gold nanorods (dimension ∼ 2 mm in length and 1 mm in width) was transferred onto the distal end of the locust wing (n = 12 locusts). To ‘tattoo’ the film onto the wing, the silk film was first transferred onto the D.I. water. This resulted in a free floating film that was manually transferred onto the locust wing by gently wicking the silk film off the water surface. This procedure resulted in successful film transfer in most cases and the film conformed to the wing surface uniformly. A random cohort of 14 locusts with no silk film tattoos were used for the control experiments. All experiments were carried out in a blind manner where the experimenter did not know whether a particular locust was tattooed or not.

Each locust was put in a T-maze assay. ^8^ Briefly, each locust was restrained in a custom design holder that was located at one end of the longer T-maze arm (44 cm in length, elevated 11 cm from the base of the arena). Initially, the outlet of the holder was kept closed with a plastic mesh. After the locust was habituated, the plastic mesh was removed and a focused laser beam (λ=808 nm) was manually focused onto the distal end of the wing. To eliminate any experimenter bias, multiple experimenters did this test not knowing which locusts were exposed to the film and which were not. A video camera (Microsoft webcam) was used to capture the behavioral responses of the locusts in the T-maze arena. The latency with which the locust started to walk out of the holder was considered as the ‘escape time,’ and was determined off-line in a double-blind manner.

